# Voxel-wise and spatial modelling of binary lesion masks: Comparison of methods with a realistic simulation framework

**DOI:** 10.1101/2021.01.11.426223

**Authors:** Petya Kindalova, Ioannis Kosmidis, Thomas E. Nichols

## Abstract

**Objectives:** White matter lesions are a very common finding on MRI in older adults and their presence increases the risk of stroke and dementia. Accurate and computationally efficient modelling methods are necessary to map the association of lesion incidence with risk factors, such as hypertension. However, there is no consensus in the brain mapping literature whether a voxel-wise modelling approach is better for binary lesion data than a more computationally intensive spatial modelling approach that accounts for voxel dependence.

**Methods:** We review three regression approaches for modelling binary lesion masks including mass-univariate probit regression modelling with either maximum likelihood estimates, or mean bias-reduced estimates, and spatial Bayesian modelling, where the regression coefficients have a conditional autoregressive model prior to account for local spatial dependence. We design a novel simulation framework of artificial lesion maps to compare the three alternative lesion mapping methods. The age effect on lesion probability estimated from a reference data set (13,680 individuals from the UK Biobank) is used to simulate a realistic voxel-wise distribution of lesions across age. To mimic the real features of lesion masks, we propose matching brain lesion summaries (total lesion volume, average lesion size and lesion count) across the reference data set and the simulated data sets. Thus, we allow for a fair comparison between the modelling approaches, under a realistic simulation setting.

**Results:** Our findings suggest that bias-reduced estimates for voxel-wise binary-response generalized linear models (GLMs) overcome the drawbacks of infinite and biased maximum likelihood estimates and scale well for large data sets because voxel-wise estimation can be performed in parallel across voxels. Contrary to the assumption of spatial dependence being key in lesion mapping, our results show that voxel-wise bias-reduction and spatial modelling result in largely similar estimates.

**Conclusion:** Bias-reduced estimates for voxel-wise GLMs are not only accurate but also computationally efficient, which will become increasingly important as more biobank-scale neuroimaging data sets become available.

## 1. Introduction

White matter hyperintensities of presumed vascular origin (WMHs), also known as white matter lesions or leukoaraiosis (Wardlaw et al., 2013), are signs of cerebral small vessel disease (SVD) in the brain. Lesions are evident on Magnetic Resonance Imaging (MRI) as hyperintensities on the T2-weighted, fluid attenuated inversion recovery (FLAIR), and proton density-weighted brain images. WMHs are common in the aging brain and are associated with cerebrovascular burden (Rostrup et al., 2012; Griffanti et al., 2018; Lampe et al., 2019). It is not clear how different contributors to the cerebrovascular burden, such as hypertension or smoking history, relate to the spatial distribution of WMHs in the brain and this has given rise to the exploitation of a variety of statistical methods in the field.

There are other types of lesions in brain imaging, which differ by the factor influencing their development. For example, multiple sclerosis (MS) is an autoimmune disease of the central nervous system, which causes the destruction of myelin further resulting in brain and spinal cord lesions. Another example are stroke lesions, which can have very similar signal intensities as WMHs and are of vascular origin too. However, independent of the type of brain lesions, their size, location, growth, etc., are important for diagnosis, treatment or prevention. While all types of lesion data motivate the current work, going forward we will focus on WMHs of presumed vascular origin.

As originally created, the MRI scans exist in so-called native space, and do not correspond to any other subject’s brain. These native space images are used to quantify the severity of lesions either based on a visual scoring system (e.g. Fazekas scale Fazekas et al. 1987), or by segmenting the lesions by producing a binary lesion map indicating lesion presence/absence. Visual scoring as well as manual lesion segmentation are quite common in neurodegenerative diseases such as MS, even though they are expensive, time-consuming and subject to inter-rater variability (Rudick et al., 2012; Hagens et al., 2019). An objective automated segmentation procedure, such as BIANCA (Griffanti et al., 2016), is preferable since it provides a scalable method to obtain reproducible lesion maps on thousands of subjects.

Whether created manually or by an automated method, the native space lesion maps can be transformed to the MNI atlas space, producing aligned binary lesion maps ready for group analyses. For example, a voxel-wise analysis can compare the distribution of patterns of lesions from different disease subtypes (Filli et al., 2012), or perform voxel-wise linear regressions between lesion probability and different clinical disability scores (Charil et al., 2003; Kincses et al., 2011). Approaches such as the ones mentioned are known as mass-univariate since they fit a model at each voxel independently, ignoring any spatial dependence between nearby voxels which is later accounted for at the inference stage (e.g. using a method like false discovery rate (FDR) correction that allows for positive spatial dependence (Genovese et al., 2002)). While some authors have used a standard linear model with lesion incidence as response (Kincses et al., 2011), this is ill-advised as it ignores the binary and heteroscedastic nature of the data.

Mass-univariate voxel-wise modelling of lesion masks that accounts for the binary nature of the data is done through maximum likelihood estimation of a generalized linear model (GLM), e.g. logistic or probit regression. While the GLM has been used in the voxel-wise brain lesion mapping literature (Lampe et al., 2019; Rostrup et al., 2012), to our knowledge the limitations of logistic or probit regression with small sample size and/or low incident responses has not been addressed. These issues have been thoroughly investigated in the statistics literature and a short overview is provided here.

Outside of linear models, maximum likelihood estimation typically requires iterative optimization, such as iteratively reweighted least squares (IRLS) (Green, 1984). When a covariate (or a combination of covariates) in a logistic or probit regression model perfectly separates the outcome variable, ‘data separation’ occurs (Albert & Anderson, 1984) and the maximum likelihood estimates (MLEs) for those covariates are infinite. Hence the iterative procedure for maximum likelihood will diverge or, even worse, stop early, reporting massive in absolute value estimates without any warning that the estimates are in reality infinite. This is more likely to happen when dealing with rare responses or small sample size. For example, with lesion data, it could happen if only subjects older than 60 years of age have a lesion at a particular voxel and no subject younger than 60 does. In such cases, estimated standard errors also diverge to infinity but faster than the estimates. As a result, the commonly used Wald statistics become artificially small in absolute value masking any significance in evidence when testing. In addition, the optimal properties of the ML estimator only hold asymptotically, and finite sample properties may be far from what is expected asymptotically. To address both limitations of the MLE, the use of a bias-reduction approach (Kosmidis & Firth, 2009; Kosmidis et al., 2020) in mass-univariate voxel-wise modelling is explored. The method guarantees finite-valued estimates (Kosmidis & Firth, 2020) as it corrects for the first-order bias of the ML estimator. Furthermore, bias-reduced estimates are fast to obtain (typically being only slightly more expensive than MLEs) and the voxel-wise modelling allows for parallel implementation, which makes the method feasible for large imaging data sets.

In contrast to the mass-univariate approaches for brain image analysis, Ge et al. (2014) introduce a Bayesian Spatial Generalized Linear Mixed Model (BSGLMM). While still accounting for the binary nature of the data, the main difference between BSGLMM and the classical GLMs is that BSGLMM accounts for the local spatial dependence in the brain through the inclusion of spatially varying coefficients to a Bayesian spatial model. Spatially varying coefficients are latent spatial processes (or fields) and they are modelled jointly using a multivariate pairwise difference prior model, a particular instance of the Multivariate Conditional Autoregressive (MCAR) model. Given that the method estimates an entire brain mask of coefficients for each covariate in a model (e.g. age and sex), there is a considerable computational burden, which is partly alleviated by a parallel graphical processing unit (GPU) implementation (Ge et al., 2014).

The motivation for the present work is the lack of validation for the mass-univariate generalized linear regression model, and the only very limited simulation framework used to evaluate the BSGLMM method. In particular, it is not known whether the sample sizes and typical incident rates found in WMH studies produce highly biased (or even divergent) estimates of regression effects with standard maximum likelihood estimators. With the Bayesian approach, while Ge et al. (2014) provide simulations showing the benefits of spatial regularization, those evaluations used only 2D images with large homogeneous lesion patterns that do not reflect the highly structured and inhomogeneous patterns found in real data.

To gain a better understating of the differences between the alternative lesion mapping methods, in this paper we develop a novel simulation framework of artificial lesion maps. We estimate the effect of age on lesion probability in a reference data set (a subset of the UK Biobank data set Miller et al. 2016) and we use it to simulate a realistic voxel-wise distribution of lesions across age. We use age as a covariate since it is thought to be the strongest risk factor for the presence of lesions, although the simulation approach could be adapted to utilise effect maps of any risk factor. To mimic the real features of lesion masks we suggest matching brain lesion summaries (total lesion volume, average lesion size and lesion count) across the reference data set and the simulated data sets. In this way, we allow for a more realistic, fairer comparison between the modelling approaches.

In this paper, we compare three alternative approaches for modelling binary lesion masks, two mass-univariate regression methods and the BSGLMM method, with the remainder of the paper organised as follows. In Section 2.1 we start by providing the details behind these different methods. We set out the steps of our proposed novel simulation framework in Section 2.2, which mimics features of real lesion masks. We then apply the three modelling approaches to simulated data sets and evaluate their performance in terms of a range of estimation accuracy metrics, such as bias and mean squared error, as well as spatial overlap between Wald statistics (reference versus estimated z-scores), false positive control and computational cost (Section 3.1). To demonstrate the scalability of one of the methods, we apply it to a subset of the UK Biobank data, where we estimate the effect of systolic blood pressure on lesion probability (Section 3.2).

Software to generate lesion masks using our simulation framework is available through the Open Science Framework website^1^, which also provides a demonstration of the parallel GLMs implementation.

## 2. Materials and Methods

### 2.1. Summary of existing regression methods

Suppose that there are *N* individuals and that each subject *i*(*i* = 1,…, *N*) comes with a binary lesion mask 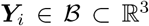. 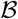 is the human brain and we consider *M* cubic cells (voxels) as a discretization of the 3D brain on a regular rectangular grid, where *s_j_* denotes the *j*th voxel within the brain (*j* = 1,…, *M*). When modelling binary lesions masks voxel-wise, we consider two approaches that ignore spatial dependence and one that explicitly models that dependence.

#### 2.1.1. Generalized Linear Model

A mass-univariate approach fits a model at each voxel marginally, ignoring spatial dependence. A Generalized Linear Model (GLM) is required in order to respect the binary nature of the data. Every GLM has a link function, deterministic and stochastic components, which we write as

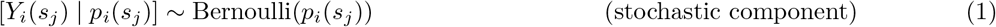

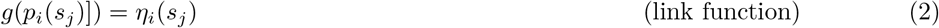

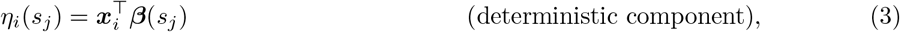

where

- *Y_i_*(*s_j_*) denotes a Bernoulli random variable with probability of success *p_i_*(*s_j_*) and probability mass function *f*(*y_i_*(*s_j_*)|***x**_i_*; ***β***(*s_j_*)), where *y_i_*(*s_j_*) is a realization of random variable *Y_i_*(*s_j_*) that represents the presence (*Y_i_*(*s_j_*)=1) or absence of a lesion for subject *i* at voxel *s_j_*. Note that in this mass-univariate voxel-wise modelling framework, *Y*_1_(*s*_1_),…, *Y_N_*(*s*_1_),…, *Y*_1_(*s_M_*),…, *Y_N_*(*s_M_*) are assumed to be independent random variables given *p_i_*(*s_j_*).
- *g* denotes the link function, which is a monotonic function that relates the expectation of the stochastic outcome to the deterministic component.
- ***x**_i_* denotes the *P*-vector of subject-specific covariates for subject *i*, where ***X*** is the full rank model matrix that collects ***x***_1_,…, ***x**_N_* in its rows and has columns ***X***_1_,…, ***X**_P_*.
- *β*(*s_j_*) = (*β*_1_(*s_j_*),…, *β_P_*(*s_j_*))^⊤^ is a *P*-vector of parameters at each voxel *s_j_*; these are fixed effects.

The GLM outlined in Equations (1–3) is fitted at each voxel *s_j_* independently. We obtain the maximum likelihood estimators (MLEs) 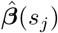 by maximizing the log-likelihood

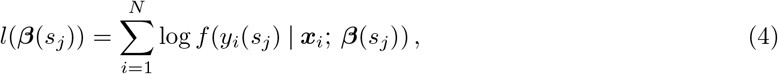

through an iterative optimization procedure, such as IRLS (Green, 1984). The MLE is typically the default choice of an estimator because of its optimal asymptotic properties (consistency, asymptotic normality and efficiency). If the model assumptions are adequate, then any inferential procedures based on those estimates, such as tests using Wald statistics (also known as standardized coefficients or z-scores) are also asymptotically correct. However, for finite sample size *N* the estimates can be unstable and biased.

Bias-reduction in parametric estimation has been thoroughly studied in the literature; for a detailed review see Kosmidis (2014). There are many methods, such as bootstrap, which correct for bias, but they rely on the existence of the MLE. However, if data separation occurs, the MLE for one or more covariates is infinite^2^, which apart from computational issues also results in invalid Wald-type inference (extremely wide and uninformative Wald-type confidence intervals due to large standard errors). The bias-correction approach which we focus on in this work was first introduced in Firth (1993) for logit link binomial GLMs and then was further developed for exponential families and applied in generalized nonlinear models (Kosmidis & Firth, 2009; Kosmidis et al., 2020). Adjustments to the score equations (partial derivatives of the log-likelihood set to zero) ensure that estimates 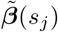 have asymptotically smaller bias than what the MLE typically has; see Appendix A.2 for details. Furthermore, obtaining the MeanBR estimates is only a modest addition to the computational complexity for computing the MLEs. The MeanBR method is implemented in the `R` package `bsglm2` (Kosmidis et al., 2020; Kosmidis, 2020) as an extension to the base `R glm tool`.

For the current analyses of simulated and real data, we have chosen probit link Φ^−1^, where Φ indicates the standard normal cumulative distribution function; we use probit link to ensure comparability with the link used in the BSGLMM approach. Finally, at each voxel, we obtain maximum likelihood estimates 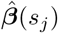 and mean bias-reduced estimates 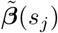 along with Wald statistics 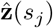 and 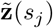 based on those estimates, respectively.

#### 2.1.2. Bayesian Spatial Generalized Linear Mixed Model

The spatial generalized linear mixed model (GLMM) is based on the GLM presented above. However, the deterministic components are explicitly defined functions of space. While the stochastic component and the link function in Equations (1) and (2) are the same, the deterministic component introduced by Ge et al. (2014) is:

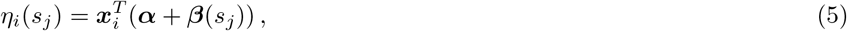

where the key difference is the inclusion of spatially varying coefficients in addition to the fixed effects. In particular,

- *α* denotes a *P*-vector of parameters, fixed effects.
- *β*(*s_j_*) denotes a P-vector of mean-zero random effects, one at each voxel *s_j_*. These random effects are spatially varying voxel-specific effects.

The last bit of the model specification is to assign priors to all parameters in order to complete the specification of the hierarchical model. This is done in the following way:

- fixed effects’ priors are flat, improper, uninformative, i.e. *π*(***α***) ∝ **1**.
- random effects (spatially varying coefficients) have Markov random field (multivariate conditional autoregressive (MCAR) model) priors to account for the spatial dependence. Two voxels are considered to be neighbors if they share a face, i.e. a maximum of six neighbors. In terms of notation, *S_k_* denotes the set of neighboring locations for location *s_j_* and the cardinality of this set is denoted as *N*(*s_j_*). The MCAR prior can be written as

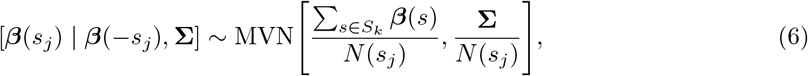

where

– ***β***(−*s_j_*) denotes ***β*** excluding the coefficients at voxel *s_j_*.
– ***β**^T^* = [***β**^T^*(*s*_1_),…, ***β**^T^*(*s_M_*)] is a *P* × *M* column vector.
– **Σ** is a *P* × *P* symmetric positive definite matrix.
– The inverse of the hyperparameter **Σ** is needed and its inverse is assumed to have Wishart prior, i.e. **Σ**^−1^ ~ W(*ν, **I**_P_*), where *ν* is set to 0 in Ge et al. (2014) and ***I**_P_* is a *P* × *P* identity matrix.

The joint distribution of ***β*** is improper and not identifiable (Ge et al., 2014), but all the full conditional distributions are well-defined but not easy to sample from. A graphical processing unit (GPU) allows for parallel implementation of the Gibbs sampler derived in Ge et al. (2014)^3^. At each voxel, posterior summaries for the spatially varying coefficients ***β***\*(*s_j_*) are obtained along with standardized posterior effects or z-scores (posterior mean divided by posterior standard deviation) **z**\*(*s_j_*).

### 2.2. Simulations

Simulation of brain lesions is complicated by the need for a generative model that accounts for dependence in the data. The mass-univariate model makes no attempt to model dependence, and while the BSGLMM explicitly models dependence, it does so on the regression parameters not the data itself. That is, the BSGLMM assumes that the binary lesion data ***Y**_i_* are independent given the (CAR-regularised) regression parameters ***β***. Thus even if an accurate regression model could be fit everywhere in the brain, simulation of lesion data ***Y*** from either the mass-univariate, or a conditionally independent Bayesian model would be characterised by independent “salt and pepper” noise, i.e. random isolated lesions of 1 or 2 voxels, or single voxel omissions from an otherwise large lesion.

Thus in this work we develop a novel simulation approach that generates realistic binary lesion data that is calibrated to a given generalized linear model. We use this approach to compare three alternative methods (see Section 2.1) for modelling the spatial distribution of white matter lesions, and to assess their performance in terms of a variety of measures of accuracy, such as mean squared error (MSE).

#### 2.2.1. Simulation procedure

We aim to simulate 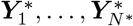 lesion masks for *N*\* subjects that follow a generalized linear model for a given map of regression parameters. Given an existing data set (referred to as ‘reference’ data) of *N* lesion masks ***Y*** = (**Y**_1_,…, ***Y**_N_*) and a vector ***X***_2_ = (*X*_12_,…, *X*_*N*2_) of centered age, artificial binary lesion masks are simulated as follows:

**Step 1:** Learn Parameters from reference data. For a reference dataset ***Y*** and a model matrix ***X*** = [**1**_*N*_ ***X***_2_], where **1**_*N*_ is an *N*-vector of ones, obtain estimated maps for intercept and age effects ***β***(*s_j_*) = [*β*_1_(*s_j_*) *β*_2_(*s_j_*)], at each *s_j_* (*j* = 1,…, *M*). These coefficients ***β*** are considered as truth going forward.
**Step 2:** Construct Simulation Design. Create the model matrix ***X***\* by simulating age 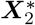 for *N*\* subjects, where 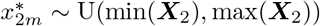, (*m* = 1,…, *N*\*), the uniform distribution on the age range in the reference data set. Then the simulation model matrix is 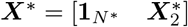.
**Step 3:** Simulate Smooth Noise for Linear Predictor. Simulate a zero-mean Gaussian Random Field (GRF) with squared exponential covariance function independently for each of the *N*\* subjects. The `R` package `RandomFields` (Schlather et al., 2020) and its function `RFsimulate()` are used to simulate a GRF with covariance *C*(*h*) = *σ*^2^ exp(−*h*^2^/2*ℓ*^2^), where *h* is the distance between voxels, and the two parameters are the variance *σ*^2^ and the scale *ℓ*. The scale determines the dependence between voxels.
**Step 4:** Generate Binary Lesion Data. Create a binary lesion mask for subject *m* as 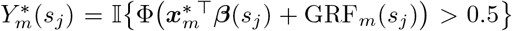, where 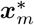 is the mth row of the simulated model matrix ***X***\* and GRF_*m*_(*s_j_*) is the value of the simulated GRF for subject *m* at voxel *s_j_*. In particular, we first add the true effect and noise and transform the sum into a lesion probability using the cumulative distribution function of the standard normal before thresholding the lesion probabilities at 0.5 to get binary lesion masks. Note that the threshold of 0.5 ensures that we match the lesion incidence found in the reference data ***Y***, set via the intercept term since we are using centered age.

In our illustration, the reference data set ***Y*** consists of binary lesion masks of 13,680 UK Biobank (UKB) (Miller et al., 2016) participants along with their age at scan date; the data set is described further in Section 2.3. Since our binary lesion mask simulator takes effect maps and GRF parameters as inputs, we make the following choices: (i) the effect maps ***β***(*s_j_*) are mean bias-reduced estimates obtained by fitting voxel-wise GLMs with probit link function and age as the only covariate in the model, and (ii) the use of probit link GLMs to model the simulated data means the variance parameter *σ*^2^ should be fixed to 1 to match the standard Normal variance, and thus there is only one free GRF parameter *ℓ*. Note that we simulate lesion masks of the same resolution as the reference data lesion masks.

#### 2.2.2. Tuning of simulation parameters

Aiming to mimic the real features of the data, we tune the scale parameter *ℓ* of the GRF to minimise the discrepancies between reference and simulated data medians of the following lesion mask summaries: (i) total lesion volume, (ii) lesion count, (iii) average lesion size. Specifically, looking over ten age bins,

i. total lesion volume is defined as the number of lesion-affected voxels;
ii. lesion count is determined using the `FSL cluster`^4^ function (connectivity 6);
iii. average lesion size is defined as total lesion volume divided by lesion count;

and the age bins are determined by the deciles of the reference data set age distribution.

We repeat Steps (3-4) on a grid of scale parameters conditionally on the noise component in Step 3 and until a satisfactory match is found between the simulated medians and reference medians across age bins.

#### 2.2.3. Measures of accuracy

Once the GRF parameter is tuned, the three regression modelling methods can be applied and their performance compared across *R* repetitions. First, we repeat Steps (3-4) (Section 2.2.1) *R* times to obtain *R* simulated data sets. Then, for each simulated data set *r*(*r* = 1,…, *R*), we fit *N*\* lesion masks ***Y***^*(*r*)^ on age 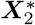 to obtain ML 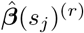, MeanBR 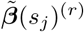 and BSGLMM 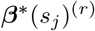 intercept and age estimated maps and their associated z-scores.

To compare the performance of the three regression modelling methods, we calculate the following measures of accuracy of MLE 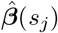, voxel-wise:

- Bias 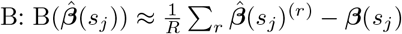,
- Mean squared error MSE: 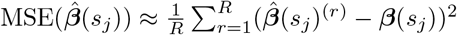,
- Probability of underestimation PU: 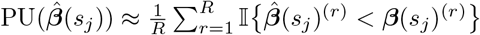.

The corresponding summaries are estimated for 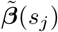 and 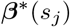. We also explore the Pearson correlation coefficient between the estimated coefficients and the reference data coefficients as another measure of estimator accuracy, resulting in one correlation coefficient per realisation *r* across the three methods.

To make inference about the effect of a covariate on the lesion probability across the brain, z-score maps are typically explored. Given the difference in the sample size *N* of the reference data set and *N*\* of the simulated data sets, the power to detect significant age effect varies and use of a fixed z-score threshold (e.g. ±1.96) to compare maps is not appropriate. Thus, we fix the z-score threshold to a particular percentile of the z-score distribution (in absolute value), such that we select the highest *M*\* z-scores. We explore the Dice similarity coefficient (DSC) (Dice, 1945) to measure the spatial overlap between a reference result and one of the three methods, e.g. the highest *M*\* reference age z-scores **z**(*s_j_*) and the the highest M* age z-scores 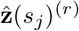 for simulated data set *r*. DSC results are the mean across *R* repetitions. We note that in the image validation literature, a DSC greater than 0.7 is interpreted as good overlap (Zijdenbos et al., 1994; Zou et al., 2004).

We also create maps of the lesion incidence across the brain, where *p*(*s_j_*) denotes the reference data set lesion incidence at voxel *s_j_* and 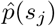 denotes the lesion incidence for a simulated data set.

Software to generate lesion masks using our simulation framework is available through the Open Science Framework website^5^, which also provides a demonstration of the parallel GLMs implementation.

### 2.3. Application

To demonstrate the scalability of the mass-univariate approaches (ML and MeanBR), we apply them to a subset of the UK Biobank data (Miller et al., 2016). The data set includes 13,680 healthy ageing individuals, for details on the selection criteria see Veldsman et al. (2020). Voxel-wise analysis is used to investigate the effect of systolic blood pressure (BP) on the spatial distribution of lesions while controlling for confounding (age, sex, age by sex interaction and head size scaling are included as confounding variables; this is the minimal set of confounding variables suggested by Alfaro-Almagro et al. 2020). The mean age of the participants is 62.9 years (±7.4 years) with 53% being female (7,236 women). Two sequential measurements of systolic BP were taken in each subject (either manual, or automatic measurement) and the average of these two readings is used as our main covariate of interest. Note that blood pressure is known to be a dominant risk factor for the presence of lesions (Debette & Markus, 2010), so it was chosen for illustrative purposes.

To generate the binary lesion masks for these 13,680 subjects, we use the Brain Intensity Abnormality Classification Algorithm (BIANCA) (Griffanti et al., 2016) to segment the lesions. BIANCA’s inputs include T1-weighted and T2-weighted FLAIR images (Alfaro-Almagro et al., 2018). The BIANCA output image in native space is thresholded at 0.8 and binarised as part of the segmentation, where the threshold is optimised as part of the BIANCA training on manually segmented masks of subjects from the UKB cohort. Those binary maps in subject space are then registered to 2mm MNI space by applying the estimated spatial normalisation parameters derived as part of the published UKB preprocessing pipeline (Alfaro-Almagro et al., 2018). More specifically, the generation of T2 FLAIR images in MNI space includes T2 FLAIR to T1 linear registration (FLIRT, Jenkinson et al. 2002) and T1 to MNI non-linear warping (FNIRT, Andersson et al. 2007). The resulting images are binarised with a 0.5 threshold, as interpolation produces non-binary values. It is these 13,680 binary lesion masks (reference data **Y**) that are fit using a mass-univariate approach (i) to define the reference MeanBR estimates for intercept and age used in the simulator (Step 1 of the simulator in Section 2.2.1 with age as the only covariate), and (ii) to obtain ML and MeanBR estimates for systolic blood pressure across voxels while accounting for confounding due to age, sex, age by sex and head size scaling.

## 3. Results

### 3.1. Results on the simulated data

The reference data set used to obtain the reference coefficients is the subset of the UKB data set described in Section 2.3. We fit 13,680 lesion masks on age to obtain MeanBR voxel-wise estimates for the intercept and age terms. The model includes only age as a covariate and the analysis mask comprises the 72,603 voxels with non-zero lesion incidence.

#### 3.1.1. Simulation setting

##### Illustration of simulation steps

To tune the GRF scale parameter, we simulate one data set of *N*\* = 1,000 subjects for various scale values. Figure 1 demonstrates the resulting simulated masks ***Y***^*^ for two scale parameter choices for subjects aged 50 and 70 years. Increasing the scale parameter increases the smoothness of the GRF (lower granularity), i.e. the scale parameter controls the number of lesions and their size; if the variance parameter is fixed, increasing the scale parameter leads to lower count but bigger size of lesions (see Figures 2 and S2). Given that the true age effect suggests higher lesion probability with increasing age, we would expect to see more lesions for an individual aged 70, which is indeed the case in the illustration in Figure 1.

**Figure 1:**
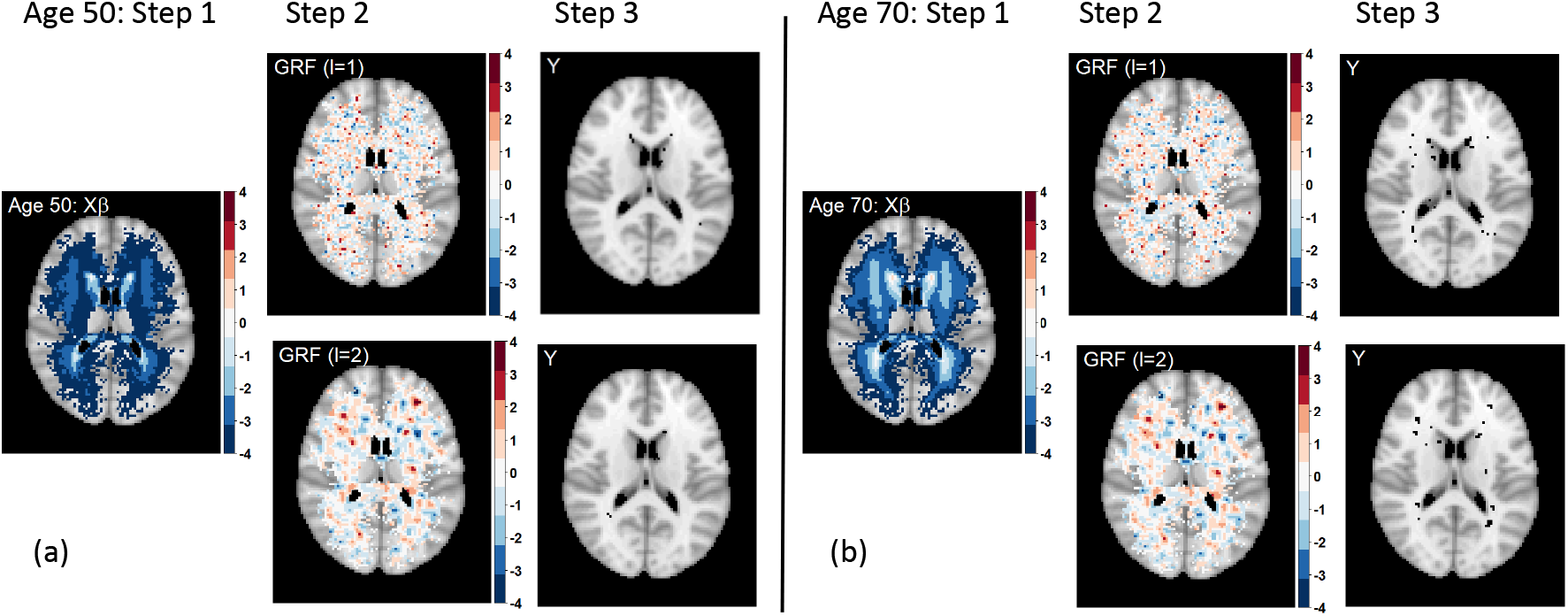
Illustration of simulation steps for scale parameters *ℓ*=1 and *ℓ*=2 for a subject aged 50 (a) and a subject aged 70 (b). Higher age is associated with higher lesion probability in the periventricular areas (Step 1), as shown by the linear predictor for a 50-year-old and a 70-year-old. Higher scale parameter leads to coarser GRF component (Step 2). The choice of the GRF parameter is crucial for simulating realistic lesion masks. Voxels with zero lesion incidence for the UKB data set (*p*(*s_j_*)=0) are plotted as transparent to show a standard anatomical MRI for reference; axial slice z=45 shown.

##### Tuning of simulation parameters

As described in Section 2.2.2, our main goal is to match as closely as possible the reference data (UKB data) lesion summaries to the simulated lesion summaries. Figure 2 includes the results for one of the lesion summaries we considered - average lesion size. The top plot represents the median average lesion size across 10 age groups for five simulation settings along with the mean and median for the reference data set (dashed and solid black lines). By visual inspection, we found that the best scale parameter value based on all three summaries (also see Figures S1 and S2) is *ℓ*=1.5. The side by side boxplots of average lesion size in one simulated data set of 1000 subjects and in the UKB data set of 13,680 participants across age groups suggests the chosen simulation setting follows closely the trend in the reference data across age and the variability in the simulated data is lower than the variability in the reference data. Note we repeated the experiment for a second seed to make sure the lesion summaries do not vary substantially and a plot complementary to Figure 2 is included in Figure S3.

**Figure 2:**
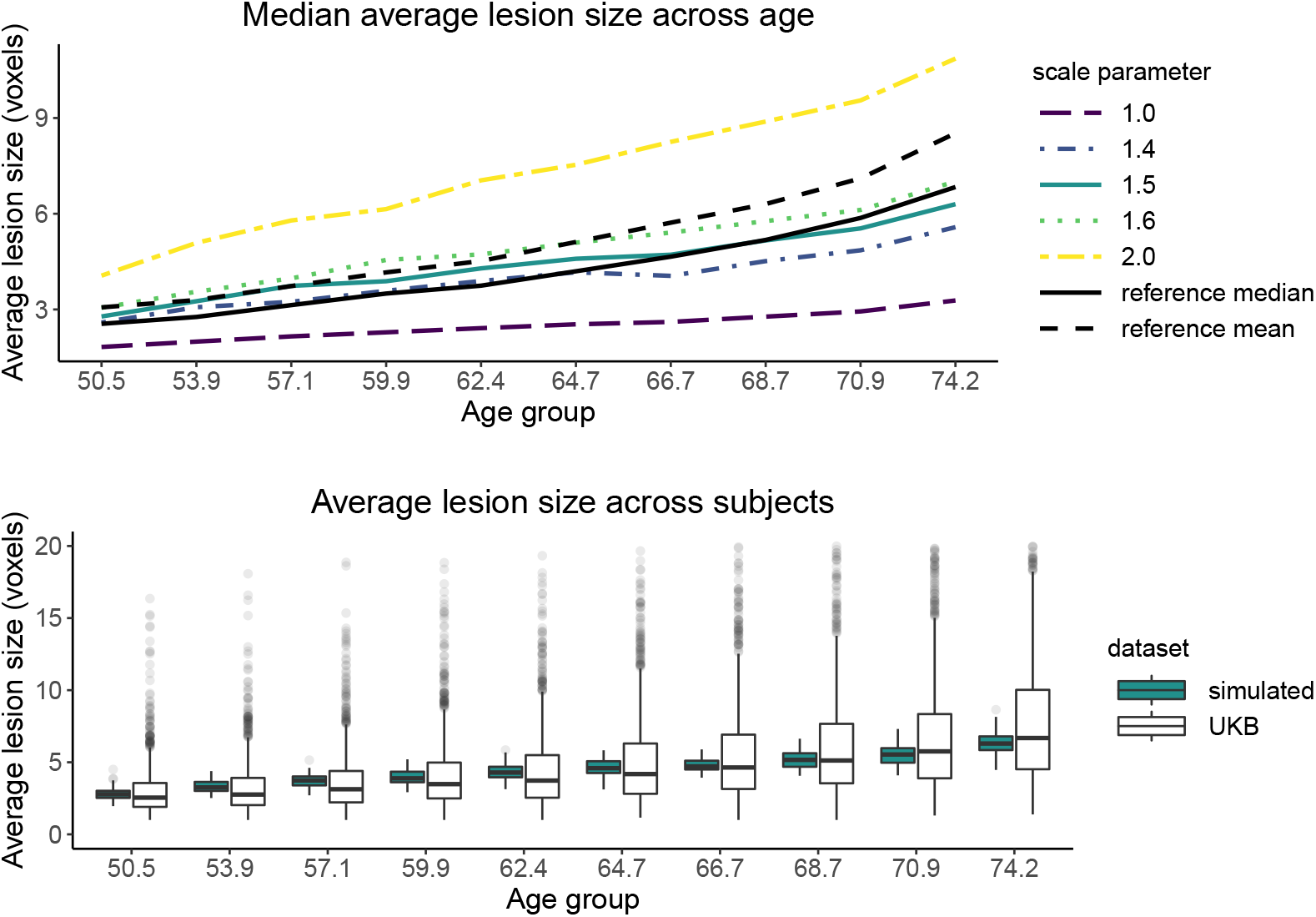
Gaussian Random Field parameter tuning by matching the reference data (UKB) median average lesion size across age bins (black solid line). (top) Plot of median average lesion size across age bins for five simulation settings (five GRF scale parameter values) and reference data values (black lines). Legend values indicate the scale parameter value *ℓ* used to simulate a GRF for each subject in the simulated sample. (bottom) Boxplots of average lesion size in UKB participants (white) and in one simulated 1000-subject sample with GRF scale parameter *ℓ*=1.5 (blue) across ten age bins. Note the x-axis labels denote the center of each age bin, the y-axis units are in 2mm^3^ voxels, and the variance GRF parameter is fixed to 1 for all simulation settings.

##### Estimated age effect

We have tuned the scale parameter of the GRF and the simulations from now on assume the variance and scale parameters are fixed to 1 and 1.5, respectively. Exploring the achieved lesion probability for a single simulated data set of 1,000 subjects and the UKB lesion probability based on 13,680 participants (Figure 3), we observe that the highest lesion probability regions are consistent across the two maps but the simulated data set does not achieve as wide a spatial coverage as the real data set (40,338 non-zero lesion incidence for the simulated data set vs 72,603 for the reference data set, respectively). Note that the UKB data set is about 14 times bigger than the simulated data set, i.e. with a single simulated data set of that size we cannot capture the rarer lesions in the outer white matter. The more limited coverage is also observed for the estimated regression coefficients since we simply do not fit the mass-univariate GLMs at voxels with zero lesion incidence. However, BSGLMM has larger z-scores due to the variance reduction of the smoothness prior and careful inspection suggests possible bleeding of signal into areas where ML and MeanBR do not capture any signal.

**Figure 3:**
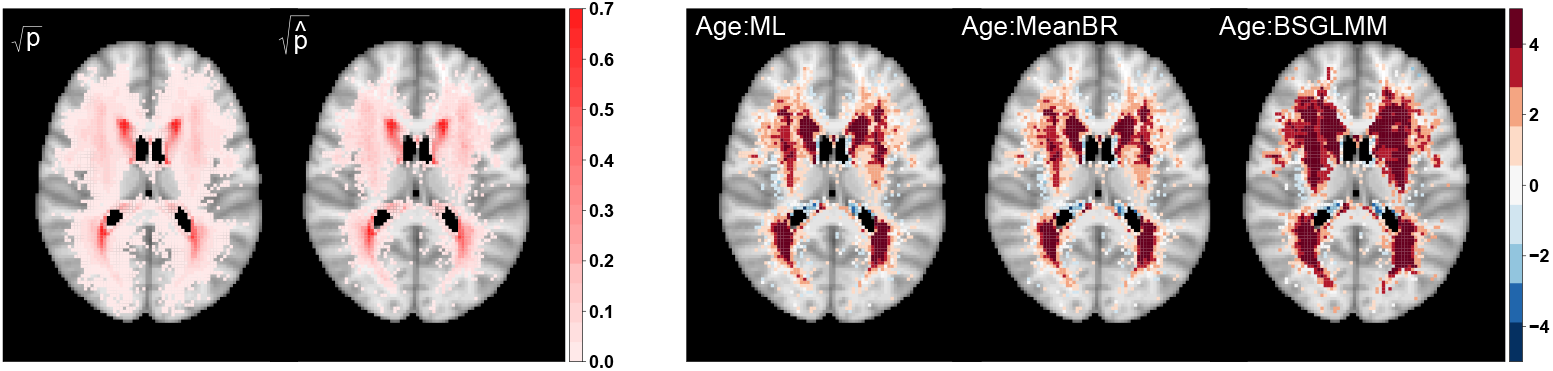
Square-root transformed lesion probability based on 13,680 UKB participants 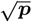 and lesion probability based on one simulated sample with *N*\* = 1,000 subjects 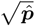 (left panel) and significance maps (z-scores) for the effect of age across methods (right panel). 72,603 voxels have non-zero lesion probability for the UKB data set and 40,338 for the simulated data set, respectively, which explains the difference in spatial coverage in the left panel.

#### 3.1.2. Estimator accuracy

We have visually compared the lesion probability maps and significance maps for one simulated data set against the UKB data set, but in order to quantify the difference between the three modelling approaches, we repeat the experiment *R*=1, 000 times, using the chosen simulation scale parameter for two sample sizes of *N*\*=250 and *N*\* = 1,000. We estimate 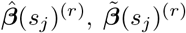 and ***β***\*(*s_j_*)^(*r*)^ (*r* = 1,…, *R*) and their associated z-scores and measures of accuracy as described in Section 2.2.3. We focus on voxels with lesion incidence in the reference data set *p* > 0.005 to ensure the lesion count is not too low in the simulated data sets.

##### Shrinkage effect

The plots of the estimated coefficients across the three methods against the reference coefficients (UKB) for age (Figures 4, S4 and S5), for one realisation of *N*\*=1,000 subjects, suggest that MeanBR and BSGLMM estimates are closer to the UKB reference coefficients than ML estimates. The plots highlight the shrinkage effect of the coefficients towards zero, especially for the voxels with the lowest lesion incidence (Figure S5). This is the result of bias reduction for MeanBR 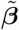 (see Kosmidis & Firth 2020) and the effect of the prior for BSGLMM ***β***^*^.

**Figure 4:**
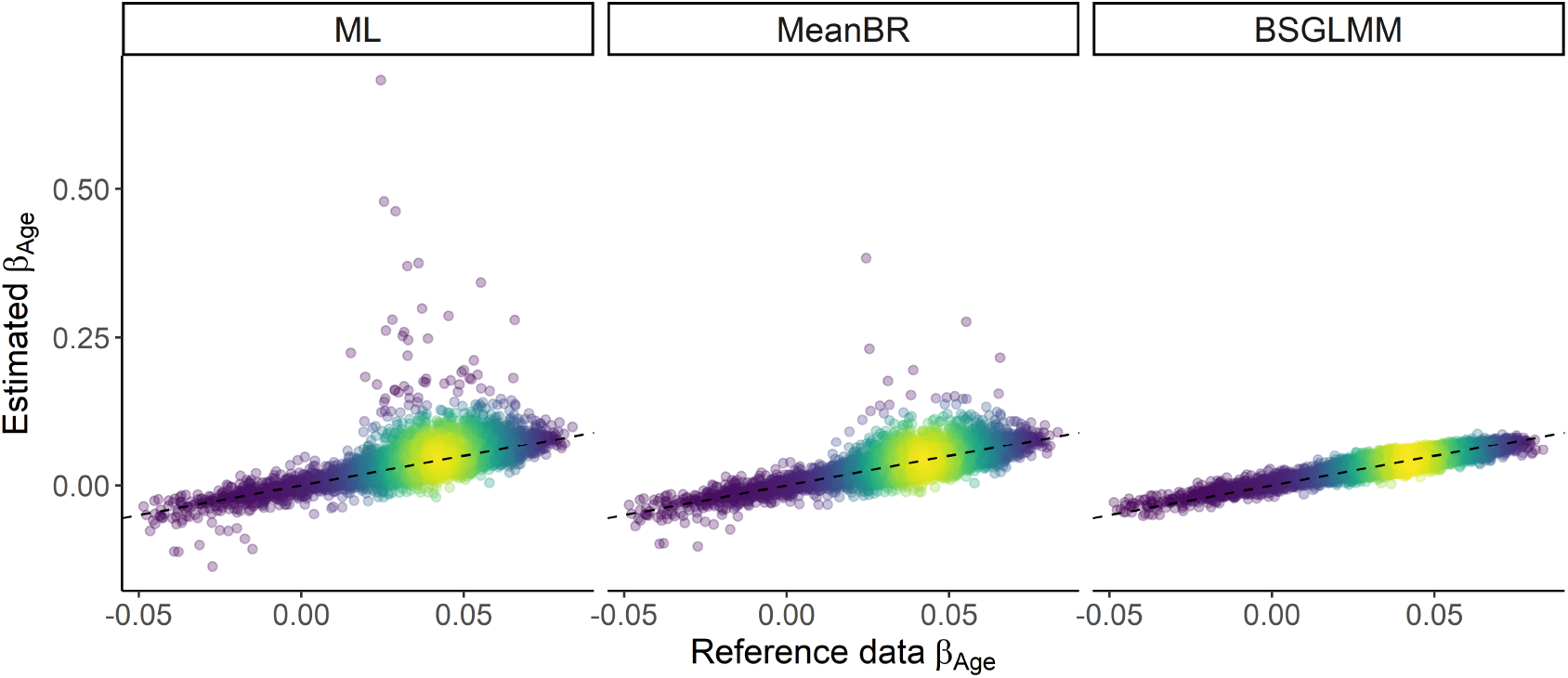
Estimated coefficients 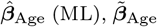 (MeanBR), 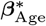 (BSGLMM) vs. ***β***_Age_ (reference). Each point is coloured according to the density of points on an invisible grid overlaid on the plots (the brighter the colour, the higher the density of the points) and the identity superimposed (dashed black line). Bias reduction and the effect of the prior result in shrinkage of the coefficients towards zero with the Bayesian model following the equality line most closely. One simulated data set of 1,000 subjects used; 11,632 voxels with reference data lesion incidence *p* > 0.005 and finite MLEs plotted.

##### Accuracy

We compare mean squared error (MSE), bias, probability of underestimation (PU) and correlation coefficient across bins of voxels for *N*\*=250 in Table 1 and for *N*\*=1, 000 in Table 2. The summaries presented for the estimates across methods are conditional on the MLEs finiteness. The Bayesian method has better performance in terms of MSE and correlation due to the smoothness prior, which reduces estimator variance at the expense of higher bias. Note that Pearson’s correlation is sensitive to outliers, thus the poor ML performance for low lesion incidence (e.g. large ML estimates as seen in Figure 4). The PU values suggest a slightly positively skewed estimates for the Bayesian method, i.e. tendency for overestimate the estimates. The opposite holds for the mass-univariate approaches, where we tend to underestimate the coefficients. According to standard asymptotic theory, all estimators should converge to a Normal distribution as the sample size increases, having PU of 50%, or equivalently being median unbiased. Reassuringly, increasing the sample size from 250 to 1000 subjects (Table 1 vs Table 2) gets PU closer to 50%. Overall, BSGLMM performs better for smaller sample size and for low lesion probability, but BSGLMM and MeanBR perform similarly for *N*\*=1, 000.

**Table 1:** Comparing methods across *R*=1, 000 data sets, *N*\*=250 subjects each. The measures of accuracy are averaged across voxel bins based on the reference data (UKB) lesion incidence *p* with standard deviation in brackets. BSGLMM is more accurate in terms of MSE and correlation values, but has higher bias than MeanBR. All values are multiplied by 1000 except probability of underestimation (PU) represented in percentage and Pearson correlation *ρ* with range (−1, 1).

| Accuracy metrics | # voxels<br>Method | $p \in (0.005, 1]$ | $p \in (0.005, 0.01]$ | $p \in (0.01, 0.05]$ | $p \in (0.05, 0.1]$ | $p > 0.1$ |
| --- | --- | --- | --- | --- | --- | --- |
|  |  | 11,634 | 4,510 | 4,981 | 935 | 1,208 |
| MSE | ML | 43.67 (49.94) | 93.63 (42.30) | 17.15 (20.66) | 0.24 (0.08) | 0.11 (0.03) |
|  | MeanBR | 1.21 (1.08) | 1.94 (1.13) | 1.01 (0.75) | 0.21 (0.05) | 0.10 (0.03) |
|  | BSGLMM | 0.21 (0.13) | 0.33 (0.13) | 0.15 (0.05) | 0.10 (0.02) | 0.07 (0.01) |
| Bias | ML | 32.12 (31.38) | 61.92 (27.75) | 18.36 (14.22) | 2.10 (1.27) | 0.81 (0.60) |
|  | MeanBR | -1.07 (2.12) | -2.75 (2.36) | -0.03 (1.12) | 0.10 (0.43) | 0.02 (0.32) |
|  | BSGLMM | 1.62 (4.94) | 2.30 (5.98) | 1.43 (4.56) | 0.92 (3.15) | 0.35 (2.03) |
| PU (%) | ML | 43.92 (4.01) | 41.25 (3.84) | 44.59 (2.94) | 47.49 (1.97) | 48.36 (1.72) |
|  | MeanBR | 57.42 (6.38) | 61.30 (7.39) | 56.34 (3.97) | 52.45 (1.89) | 51.25 (1.65) |
|  | BSGLMM | 43.97 (16.63) | 41.47 (19.47) | 44.54 (15.50) | 46.75 (12.16) | 48.78 (9.53) |
| $\rho$ | ML | 0.12 (0.013) | 0.12 (0.018) | 0.18 (0.033) | 0.79 (0.014) | 0.86 (0.009) |
|  | MeanBR | 0.46 (0.029) | 0.34 (0.034) | 0.49 (0.040) | 0.80 (0.013) | 0.86 (0.009) |
|  | BSGLMM | 0.80 (0.006) | 0.75 (0.010) | 0.81 (0.008) | 0.87 (0.009) | 0.89 (0.007) |

**Table 2:** Comparing methods across *R*=1,000 data sets, *N*\* = 1,000 subjects each. The measures of accuracy are averaged across voxel bins based on the reference data (UKB) lesion incidence p with standard errors in brackets. All values are multiplied by 1000 except probability of underestimation (PU) represented in percentage and Pearson correlation *ρ* with range (−1, 1).

| Accuracy metrics | # voxels<br>Method | $p \in (0.005, 1]$ | $p \in (0.005, 0.01]$ | $p \in (0.01, 0.05]$ | $p \in (0.05, 0.1]$ | $p > 0.1$ |
| --- | --- | --- | --- | --- | --- | --- |
|  |  | 11,634 | 4,510 | 4,981 | 935 | 1,208 |
| MSE | ML | 1.61 (14.45) | 3.89 (22.76) | 0.23 (3.36) | 0.05 (0.01) | 0.03 (0.01) |
|  | MeanBR | 0.22 (0.31) | 0.40 (0.44) | 0.15 (0.06) | 0.05 (0.01) | 0.03 (0.01) |
|  | BSGLMM | 0.07 (0.03) | 0.08 (0.03) | 0.06 (0.02) | 0.04 (0.01) | 0.02 (0.01) |
| Bias | ML | 3.13 (3.32) | 5.87 (3.68) | 1.85 (1.24) | 0.46 (0.35) | 0.21 (0.20) |
|  | MeanBR | 0.16 (0.50) | 0.31 (0.65) | 0.08 (0.39) | 0.01 (0.22) | 0.01 (0.15) |
|  | BSGLMM | 0.80 (2.95) | 1.13 (3.86) | 0.75 (2.49) | 0.31 (1.34) | 0.13 (0.81) |
| PU (%) | ML | 47.34 (2.23) | 46.27 (2.25) | 47.61 (1.90) | 48.83 (1.74) | 49.10 (1.59) |
|  | MeanBR | 53.22 (2.68) | 54.79 (2.68) | 52.83 (2.08) | 51.27(1.61) | 50.53 (1.56) |
|  | BSGLMM | 46.24 (13.29) | 44.75 (16.21) | 46.48 (11.96) | 48.41 (8.91) | 49.15 (6.96) |
| $\rho$ | ML | 0.58 (0.144) | 0.45 (0.134) | 0.81 (0.029) | 0.94 (0.003) | 0.96 (0.002) |
|  | MeanBR | 0.76 (0.022) | 0.65 (0.032) | 0.83 (0.006) | 0.94 (0.003) | 0.96 (0.002) |
|  | BSGLMM | 0.90 (0.002) | 0.85 (0.004) | 0.90 (0.003) | 0.95 (0.003) | 0.96 (0.002) |

If we further explore the spatial overlap between the highest *M*\* voxels, the DSCs across methods suggest good spatial overlap (Table 3). If we select a small number of voxels, i.e. voxels with the highest M*=1,000 z-scores, all methods seem to detect the strongest age effect very well. The Bayesian method has the worst overlap between the three methods. We understand this to be a reflection of the BSGLMM’s tendency to “bleed out” stronger effects into weaker effect areas, a problem perhaps more severe at *N*\* =250.

**Table 3:** Dice similarity coefficient (DSC) when comparing reference (UKB) and simulation z-scores estimated across the three regression methods. DSCs are obtained across *R*=1,000 data sets, *N*\* ∈ {250,1000} subjects each and the spatial overlap considered is between the highest *M*\* z-scores, where *M*\* ∈ {1000, 5000, 10000}.

| $N^*$ | Method | Dice similarity coefficient | | |
| --- | --- | --- | --- | --- |
| | | $M^*=1,000$ | $M^*=5,000$ | $M^*=10,000$ |
| $N^* = 250$ | ML | 0.824 | 0.774 | 0.748 |
|  | MeanBR | 0.821 | 0.756 | 0.718 |
|  | BSGLMM | 0.740 | 0.704 | 0.736 |
| $N^* = 1000$ | ML | 0.907 | 0.871 | 0.857 |
|  | MeanBR | 0.907 | 0.868 | 0.848 |
|  | BSGLMM | 0.888 | 0.813 | 0.815 |

##### False positive control

To further compare the methods in terms of false positive detection, we simulate a single data set with no age effect added to the true effect component in Steps 1 and 2, Section 2.2.1 (only reference data (UKB) intercept map used), but we add age 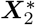 as a covariate when fitting all three models. We explore the same accuracy metrics as in Tables 1 and 2 by setting ***β***(*s_j_*) = 0 for all voxels *s_j_* and the Bayesian method performs best in terms of lowest MSE and the percentage of underestimation is very close to 50%, which is to be expected if the estimates are symmetric around zero (Table 4). Interestingly, the effect of the prior in the Bayesian method (shrinkage towards zero) reduces the bias to be smaller or comparable to the MeanBR method since the true effect is set to zero in this case. We observe (Table 5) that all methods appear to be conservative in their false positive rate especially for low lesion incidence voxels with the Bayesian method always being most conservative. This is also evident from the quantile-quantile plots (see Figure S6), where as lesion incidence decreases, the normality deviations increase.

**Table 4:**
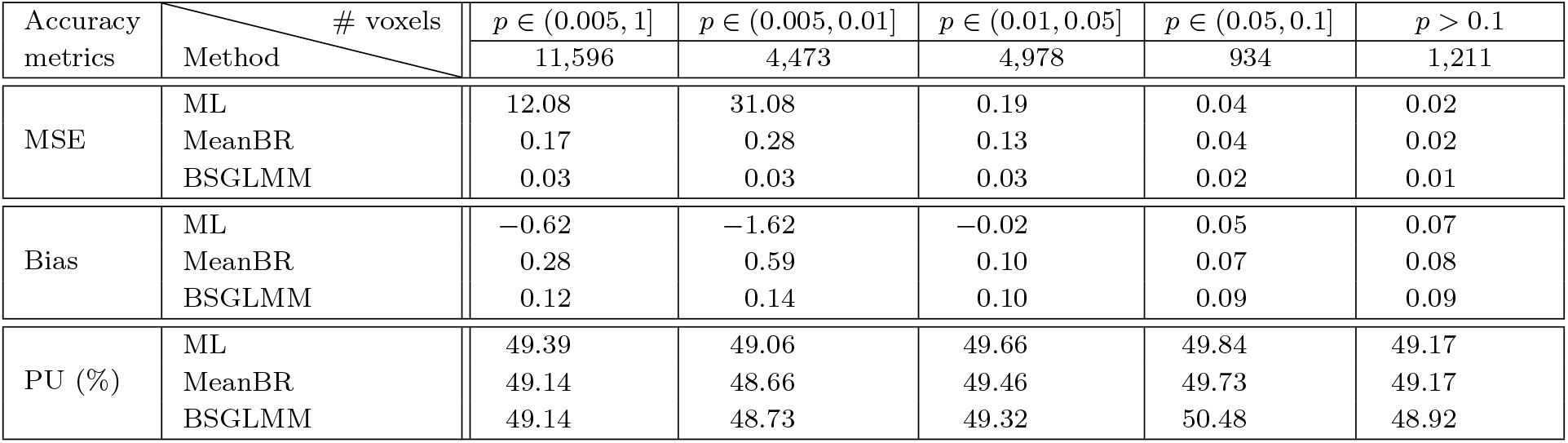
Comparing methods across one null age effect data set of *N*\* =1, 000 subjects. The measures of accuracy are averaged across voxel bins based on the reference data (UKB) lesion incidence p. All values are multiplied by 1000 except probability of underestimation (PU) represented in percentage.

| Accuracy metrics | # voxels<br>Method | $p \in (0.005, 1]$ | $p \in (0.005, 0.01]$ | $p \in (0.01, 0.05]$ | $p \in (0.05, 0.1]$ | $p > 0.1$ |
| --- | --- | --- | --- | --- | --- | --- |
|  |  | 11,596 | 4,473 | 4,978 | 934 | 1,211 |
| MSE | ML | 12.08 | 31.08 | 0.19 | 0.04 | 0.02 |
|  | MeanBR | 0.17 | 0.28 | 0.13 | 0.04 | 0.02 |
|  | BSGLMM | 0.03 | 0.03 | 0.03 | 0.02 | 0.01 |
| Bias | ML | -0.62 | -1.62 | -0.02 | 0.05 | 0.07 |
|  | MeanBR | 0.28 | 0.59 | 0.10 | 0.07 | 0.08 |
|  | BSGLMM | 0.12 | 0.14 | 0.10 | 0.09 | 0.09 |
| PU (%) | ML | 49.39 | 49.06 | 49.66 | 49.84 | 49.17 |
|  | MeanBR | 49.14 | 48.66 | 49.46 | 49.73 | 49.17 |
|  | BSGLMM | 49.14 | 48.73 | 49.32 | 50.48 | 48.92 |

**Table 5:** False positive rate evaluation. Number of voxels with z-scores significant at 5% for a two-sided test for age when no age effect is included. Percentage is calculated within each bin (column) based on the reference data (UKB) lesion incidence p. All methods appear to be conservative since we would expect 5% false positives, i.e. about 700 voxels across each row. These results are based on one simulated data set of 1,000 subjects used, 11,596 voxels with lesion incidence in the reference data greater than 0.005 and infinite MLEs discarded. Note, for this one simulated data set a 5% FDR correction found no significant voxels for any method.

| Method | $N(\vert \mathbf{z}\vert > 1.96)$ | % voxels (# voxels) | | | | |
| --- | --- | --- | --- | --- | --- | --- |
| | | $p \in (0.005, 1]$ | $p \in (0.005, 0.01]$ | $p \in (0.01, 0.05]$ | $p \in (0.05, 0.1]$ | $p > 0.1$ |
|  |  | 11,596 | 4,473 | 4,978 | 934 | 1,211 |
| ML |  | 3.6% (422) | 2.1% (95) | 4.5% (225) | 5.5% (51) | 4.2% (51) |
| MeanBR |  | 3.5% (406) | 2.3% (104) | 4.1% (205) | 5.2% (49) | 4.0% (48) |
| BSGLMM |  | 1.6% (182) | 0.4% (17) | 1.8% (88) | 3.6% (34) | 3.6% (43) |

#### 3.1.3. Computational time and scalability

On average, ML and MeanBR take about 15-20 minutes for each 250-subject data set and about 50-60 minutes for each 1,000-subject data set for single-core jobs, and 3-4 minutes and 10-12 minutes for parallel jobs (8 cores), respectively. The difference in computational cost between ML and MeanBR is minimal with bias-reduction increasing the computational cost by only a few minutes for a 1000-subject data set. Note that the number of regressions per simulated data set varies depending on the number of non-zero lesion incidence voxels. 23,404 regressions are performed on average per 250-subject data set and 40,286 per 1000-subject data set, respectively, instead of 228,483 (voxels in the brain mask). Our code determines the voxels with non-zero lesion incidence first and creates a matrix of binary values only for those voxels to be used as input to the GLMs. This trick saves computation time, but also allows better RAM management for big UKB-scale data sets since it avoids reading in all lesion masks at once. For the simulated data sets this implementation might not be optimal in terms of speed, but it makes the UKB application possible even without parallel implementation.

BSGLMM takes about 16 minutes for 100,000 iterations of the Gibbs sampler for a 250-subject data set and about 60 minutes for a 1,000-subject data set, respectively. BSGLMM is performed on an NVIDIA TESLA K80 GPU card with 12 GB RAM and 2,496 threads. While the BSGLMM run time is comparable to the ML and MeanBR, note that there is a practical upper limit of subjects due to a GPU RAM constraint; the problem arises since the Bayesian method implementation loads all binary masks limiting its application to UKB-scale data.

To summarise, while BSGLMM’s GPU implementation is computationally efficient, the ML and MeanBR have more flexibility in how parallelism can be used, making the latter easier to apply at biobank scale.

### 3.2. Results on the real data

We choose to fit the mass-univariate voxel-wise GLM with MeanBR estimates due to its scalability to the UK Biobank data set of 13,680 subjects, but also obtain MLEs to check how often separation occurs. The models we fit include systolic BP as the main effect of interest and age, sex, age by sex interaction and head size scaling as confounders. On 72,603 regressions across the brain (voxels with non-zero lesion incidence), sex MLEs are infinite for 23,330 voxels (32%); separation is more likely to occur for binary covariates and, for example, systolic BP MLEs are infinite for only 260 voxels (0.4%).

#### Spatial distribution of lesions

The lesion incidence across 13,680 UKB participants suggests the areas with the highest probabilities cover the periventricular and deep white matter regions (Figure 5). Fitting voxelwise GLMs with systolic BP as our main covariate of interest, we explore its effect on lesion probability (Figures 5 and S7). Figure 5 includes axial slices of z-scores for the effect of systolic BP (right) along with the UKB lesion probability (left); the darker the colour, the stronger the effect of systolic BP on lesion probability. The spatial distribution of lesions mirrors what is well known clinically, that is lesions are classically found capping the ventricles, clustering around the ventricles and within the deep white matter (Fazekas et al., 1987). Hypertension is known to be one of the strongest predictors of the presence of lesions (Dufouil et al., 2001). Consistent with the literature, we find hypertension related lesions distributed in periventricular and deep white matter regions as well as capping the ventricles (Moroni et al., 2018). We get 14,108 voxels with z-scores greater than 1.96 in absolute value (in comparison, 11,251 for z-scores based on MLEs, respectively). Thus, systolic BP has a strong effect on lesion probability as expected based on the existing literature.

**Figure 5:**
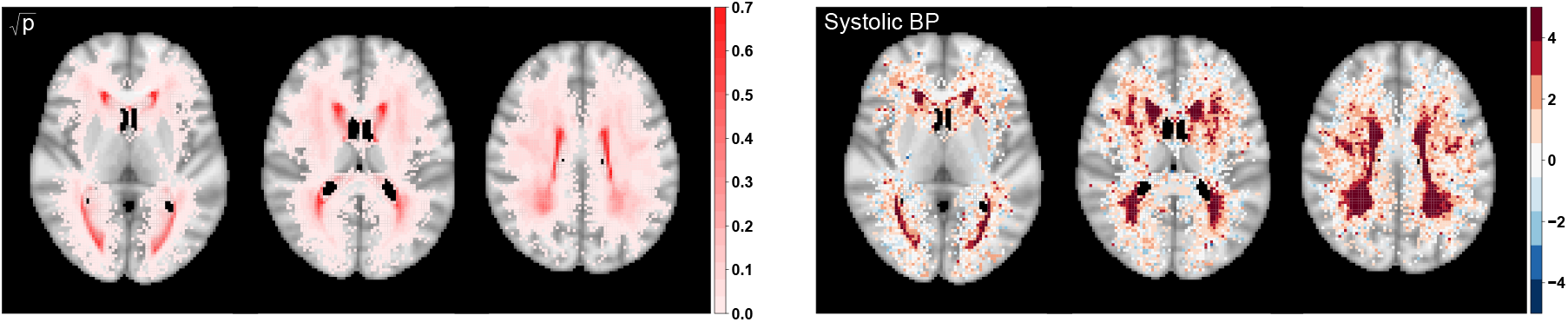
Square-root transformed lesion probability based on 13,680 UKB participants 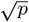 (left panel) and significance maps (z-scores based on MeanBR estimates) for the effect of systolic BP (right panel); age, sex, age by sex interaction and head size scaling included as confounders. 72,603 voxels have non-zero lesion probability; axial slices {40,45, 50}.

#### Computational time

ML and MeanBR take about 3 hours and 3.5 hours, respectively, utilizing batch jobs run on 8 cores. As mentioned in Section 3.1.3, we use the empirical incidence mask to select the non-zero incidence voxels and then run the GLMs for those voxels only, here 72,603 regressions.

## 4. Discussion

### Simulation framework

Using binary lesion masks of 13,680 healthy aging UK Biobank participants as our reference data set, we develop a binary lesion mask simulator. Age is used as the sole regressor and by using a reference data set, we start building our simulation study by setting the true age coefficient map to the UKB-derived one. In other simulation studies, binary lesion masks or 2D slices are simulated, but the true coefficients are not available (Sundaresan et al., 2019; Ge et al., 2014), which does not allow any comparison between competing methods.

We made the artificial lesion masks as realistic as possible through tuning to make sure the artificial and real lesion masks share important lesion characteristics, such as lesion size, lesion count and lesion volume. This step potentially overcomes the drawbacks of other simulation approaches, where the same count, size and shape lesions are simulated (6 spheroid lesions of size 5 voxels in each dimension Chard et al. 2010) or smoothing of the resulting simulated lesion masks (Sundaresan et al., 2019) is applied, which could introduce stronger spatial dependencies than what is expected from real lesion masks.

Our simulator code is available^6^ and ready to use to simulate binary lesion masks for healthy aging individuals. However, if the simulation framework is to be adjusted to any patient reference data set, e.g. Dementia patients, binary lesion masks and age for those patients are needed to obtain the coefficient maps and to tune the simulator as described in Section 2.2.2.

### Method comparison

We compare three alternative regression approaches for modelling of binary lesion masks. Two of them rely on voxel-wise fitting of generalized linear models using maximum likelihood and mean bias-reduction. The other is a Bayesian hierarchical model that takes into account the spatial dependence in the brain though the inclusion of spatially varying coefficients.

The bias and mean squared error of the maximum likelihood and mean bias-reduced coefficients suggest poorer performance of the maximum likelihood estimator, which is in line with the widely dispersed MLEs in the coefficient plots (Figure 4). BSGLMM seems to perform slightly better in terms of mean-squared error values, but has higher bias than mean bias-reduced estimates due to the spatial regularization it imposes. When comparing the ability of the methods to detect the voxels with the strongest age effect on lesion probability, all three methods seem to perform similarly well. Null simulations find that for a non-existent age effect all methods are valid but conservative, with false positive rates lowest for low incidence voxels.

The resources required to apply those methods vary significantly since the Bayesian spatial model utilises a GPU implementation to decrease the computational burden. For the size of the simulated data sets, all methods are relatively fast to perform with about an hour run-time for one data set of sample size 1,000 subjects (single core for mass-univariate). However, for the spatial model there is a practical upper limit on the number of subjects due to the GPU RAM constraint since all lesion masks need to be loaded in memory. On the contrary, for the mass-univariate methods, parallel implementation is possible given that the methods are applied independently at each voxel. Thus, mass-univariate approaches are computationally practical for large data sets.

### UK Biobank application

Reassuringly, the distribution of lesions in the real data reflects the known distribution of lesions associated with age and hypertension (Dufouil et al., 2001). Further work by our group demonstrates the clinical utility of the mass-univariate method (mean bias-reduced estimates) in mapping the spatial distribution of lesions associated with different cerebrovascular risk factors (Veldsman et al., 2020). Application of the mass-univariate methods (ML and MeanBR) to lesion masks on 13,680 subjects demonstrates that total separation occurs quite often for binary covariates (32% of voxels have infinite sex estimates) even in such big data sets, thus mean bias-reduced estimates would be favoured. The run-time of about 6 hours suggests that voxel-wise modelling is feasible for large data sets; heavier parallelism (we use a maximum of 8 cores) can reduce run-time substantially.

### Limitations

Our simulation framework is not adapted for automated tuning, i.e. a grid of scale values for the Gaussian Random Field are explored. An automated procedure could be developed but the merits might not outweigh the computational effort. Further improvement could be introduced by allowing the GRF scale parameter to vary across age groups to achieve a closer match to the suggested empirical lesion summaries. To match the variability in the reference data better, a more flexible covariance function than the squared exponential (e.g. Matern at the expense of an extra parameter to tune) or a non-stationary GRF might need to be adopted. However, our goal is to provide a simulation framework for the comparison of lesion mapping methods and we believe that matching the median lesion summaries across age groups is sufficient for the fair comparison of the three approaches and any alternatives that may result from future research.

Note that we do not account for any left-right symmetry of lesions, we have not imposed any physiological boundaries or 3D dependence in the entire brain when simulating the lesion masks. However, we do not believe this has any impact on the results presented here since the mass-univariate approaches do not account for the spatial dependence in the brain and the spatial model only accounts for local spatial dependence. We refer to the lesion masks as ‘realistic’ but this is not meant to imply any clinical realism (given the drawbacks mentioned) and we see the lesion masks as useful in a methods development or methods comparison context.

We generate the lesion masks in MNI space by using outputs from the published UK Biobank pipeline (Alfaro-Almagro et al., 2018). Sensitivity analysis to registration or lesion segmentation approaches could be of future interest but it is out of the scope of the current statistical work since we focus on masks in MNI space for the design of the simulation framework. The proposed lesion mask simulator could be tuned to reflect features of lesions independent of the image resolution, but the method comparison results presented are specific to the sampling resolution of 2mm^3^ voxels and we have not performed sensitivity analysis to other voxel sizes.

Note that the lack of scalability of the Bayesian approach is due to the GPU memory constraint and it could be overcome by either a time-consuming CPU implementation of the Gibbs sampler proposed by Ge et al. (2014), or by adopting a divide-and-conquer method for Bayesian inference. The latter involves splitting the data into smaller subsets (computationally manageable), sampling from the posterior distribution on all subsets and then combining the posterior samples to approximate the full data posterior, where possible methods include the ones suggested by Srivastava et al. (2018); Minsker et al. (2017); Li et al. (2017). We have focused our method comparison on the implementation available instead.

Investigating the effect of systolic blood pressure on lesion probability, we present test statistics at all non-zero lesion incidence voxels to demonstrate the scalability of the method. We could have excluded voxels where the lesion incidence fell too low and then use false discovery rate correction to account for multiple testing (Veldsman et al., 2020) to achieve better inference.

### Conclusion

The proposed simulation framework mimics real features of the data, which allows for a fair comparison between the lesion mapping methods through realistic experiments. Our findings suggest that bias-reduced estimates for voxel-wise binary-response generalized linear models overcome the instabilities of maximum likelihood estimates, and scale well for large data sets due to parallel implementation. Contrary to the assumption of spatial dependence being key in lesion mapping, our results show that voxel-wise bias-reduction and spatial modelling result in largely similar estimates, but bias-reduction is computationally feasible for biobank-scale neuroimaging data.

## Acknowledgements

Thank you to Timothy Johnson for his valuable input on the testing of our simulation framework. We would also like to thank Michele Veldsman for her help with the clinical interpretations of the real data analysis.

## Appendix A. Iterative estimation: maximum likelihood and bias-reduction

### Appendix A.1. Maximum likelihood estimates

The typical iterative algorithm used to find the maximum likelihood estimates (MLEs) for generalized linear models (GLMs) is iteratively reweighted least squares (IRLS) (Green, 1984). IRLS is equivalent to Fisher scoring obtain an iterative solution to the estimating equations (also known as score equations)

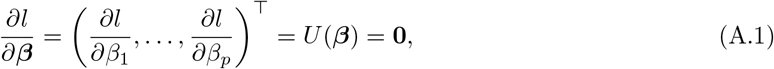

where *l* is the log-likelihood, *U*(***β***) is the score vector (*p*-vector). A Taylor series expansion for *∂l/∂**β*** (Equation (A.1)) gives the standard Newton-Raphson method for solving the estimating equations

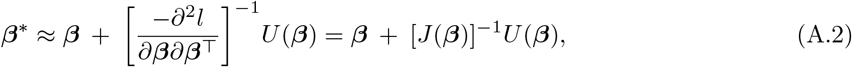

where *J*(***β***) = −*∂*^2^*l*/*∂**β**∂**β***^⊤^ is the observed information matrix, ***β*** is the initial value of the parameters and ***β***\* is the updated value. Evaluation of *U* and *J* is repeated until convergence and the resulting estimates are the MLEs we report in the paper denoted as 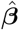.

If we replace the observed information *J*(***β***) with the expected information (Fisher information) 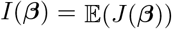 in the Newton-Raphson, the Fisher scoring iteration results. Fisher scoring is typically preferred since the Fisher information is useful post-hoc to estimate the asymptotic variance of the parameters. Note that for canonical link (e.g. logit link function for Binomial GLMs), observed and expected information coincide, hence Fisher scoring is equivalent to Newton-Raphson.

### Appendix A.2. Bias-reduced estimates

The bias-correction method we use to obtain mean bias-reduced (MeanBR) estimates 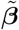 was first introduced in Firth (1993) and was then applied and developed further for exponential family models (Kosmidis & Firth, 2009; Kosmidis et al., 2020). The method is known as adjusted score equations, i.e. a penalty *A*(***β***) is added to the score equations in Equation (A.1) in order to get estimates with asymptotically smaller bias

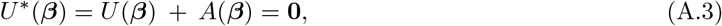

where *A*(***β***) is a p-vector based on the expected information matrix *I*(***β***) and on the observed information *J*(***β***). General formulae for the adjusted score equations are derived by Kosmidis & Firth (2009), showing that solving the mean bias-reducing score functions by iterative optimization (e.g. IRLS) results in higher-order mean unbiased estimators. What is interesting is that the general form of the first order bias is of the form

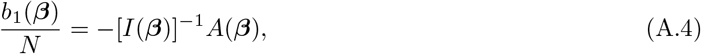

where the mean bias function *B*(***β***) of the MLE of ***β*** can be expanded in decreasing powers of *N* as

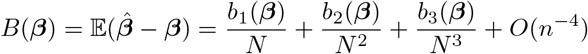

for an appropriate set of functions *b*_1_(***β***), *b*_2_(***β***),…, which are *O*(1) as *n* → ∞. Thus, the adjustment to the score functions *A*(***β***) is a function of the first-order bias and the Fisher information, i.e. iteratively subtracting the first-order bias in the Fisher scoring updates (Kosmidis & Firth, 2010). The iterative procedure from Equation (A.2) becomes a quasi Fisher scoring to obtain MeanBR estimates

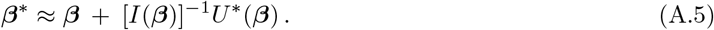

Here it is ‘quasi’ since we are using the expectation of the second derivatives of the scores *U*(***β***), instead of the second derivative of the adjusted scores *U*\*(***β***). Note that the iterated first-order bias adjustment is only possible when *b*_1_(***β***) is available in closed-form.

## Supplementary material

**Figure S1:**
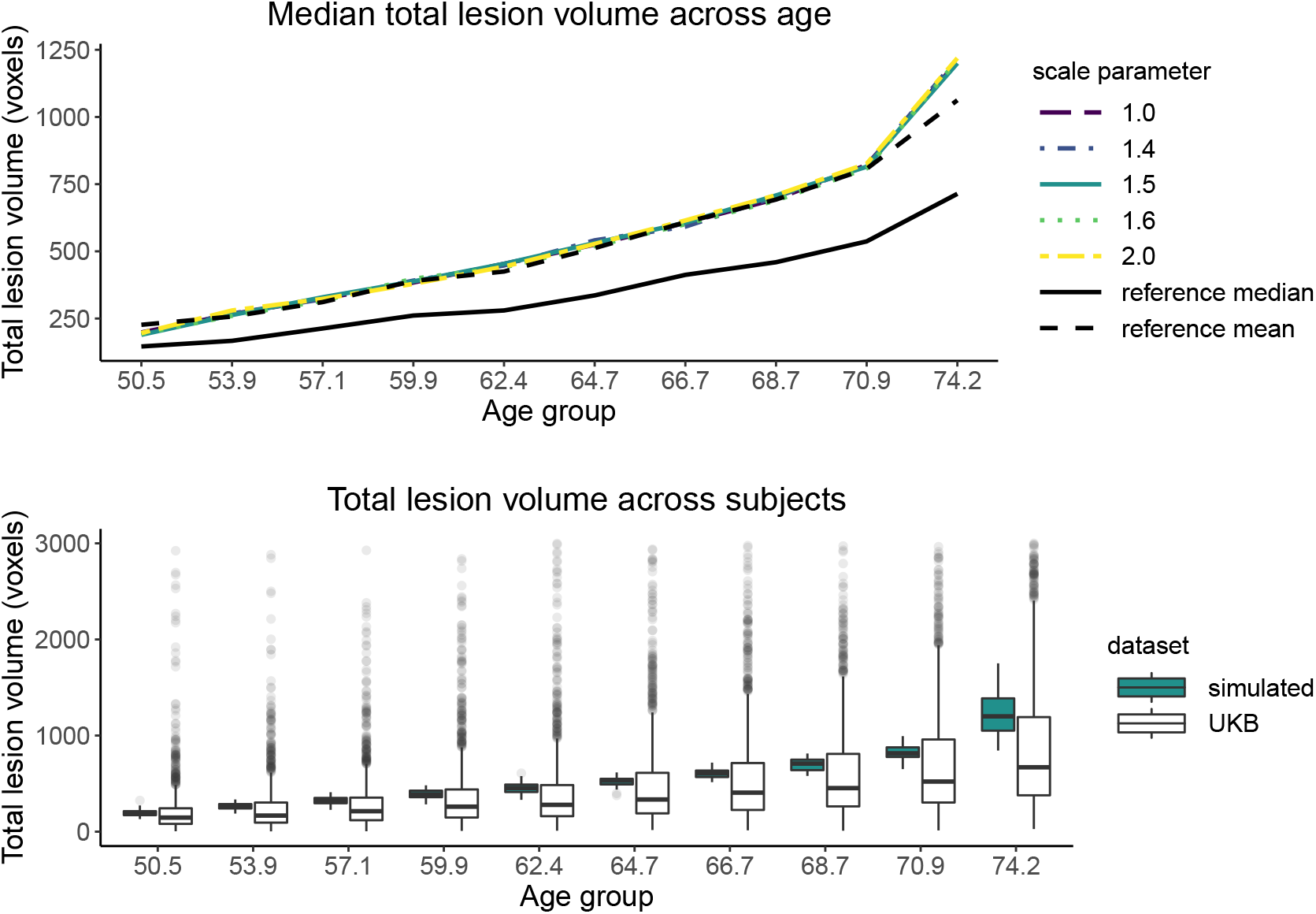
Gaussian Random Field parameter tuning by matching the reference data (UKB) median total lesion volume across age bins (black solid line). (top) Plot of median total lesion volume across age bins for five simulation settings (five GRF scale parameter values) and reference data values (black lines). Legend values indicate the scale parameter value *ℓ* used to simulate a GRF for each subject in the simulated sample. (bottom) Boxplots of total lesion volume in UKB participants (white) and in one simulated 1000-subject sample with GRF scale parameter *ℓ*=1.5 (blue) across ten age bins. Note the x-axis labels denote the center of each age bin, the y-axis units are in 2mm^3^ voxels, and the variance GRF parameter is fixed to 1 for all simulation settings.

**Figure S2:**
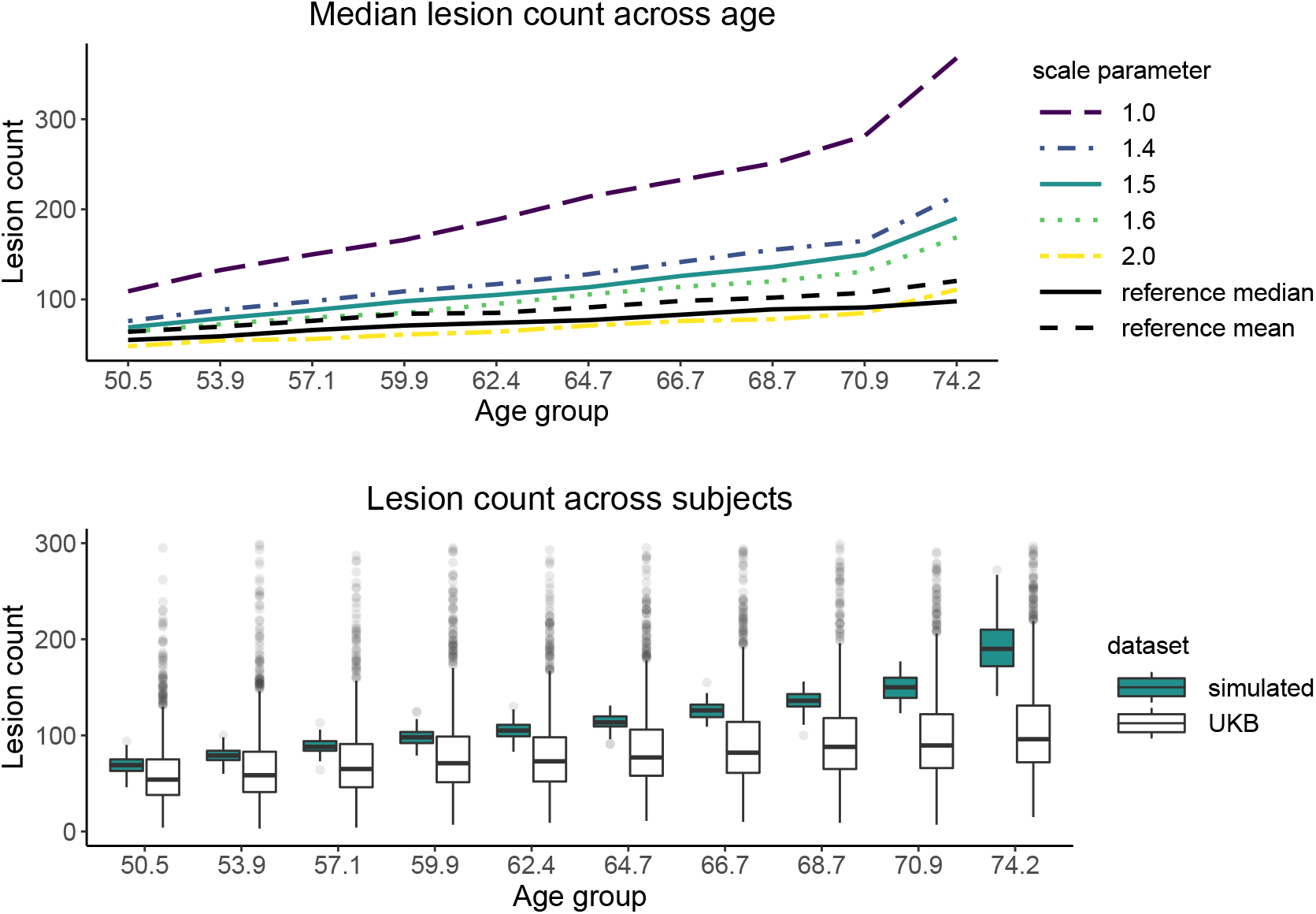
Gaussian Random Field parameter tuning by matching the reference data (UKB) median lesion count across age bins (black solid line). (top) Plot of median lesion count across age bins for five simulation settings (five GRF scale parameter values) and reference data values (black lines). Legend values indicate the scale parameter value *ℓ* used to simulate a GRF for each subject in the simulated sample. (bottom) Boxplots of lesion count in UKB participants (white) and in one simulated 1000-subject sample with GRF scale parameter *ℓ*=1.5 (blue) across ten age bins. Note the x-axis labels denote the center of each age bin, the y-axis units are in number of connected components (lesions), and the variance GRF parameter is fixed to 1 for all simulation settings.

**Figure S3:**
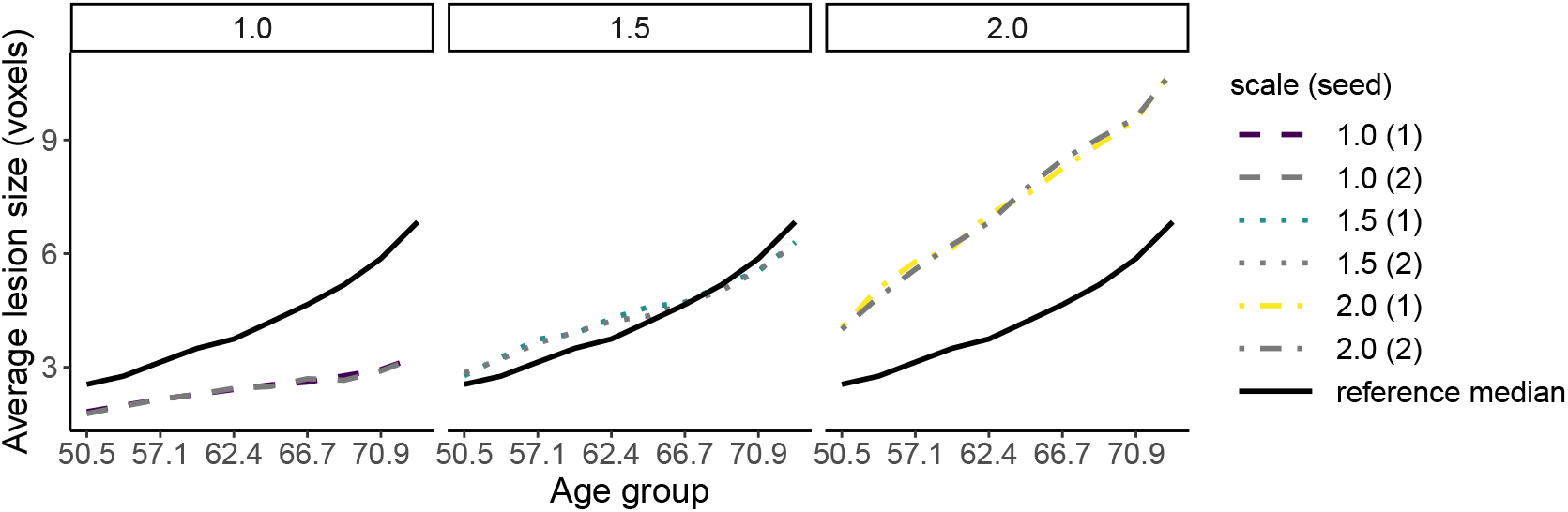
Gaussian Random Field parameter tuning by matching the reference data (UKB) median average lesion size across age bins (black solid line) replicated for two seeds. Legend values indicate the scale parameter value *ℓ* used to simulate a GRF for each subject in the simulated sample and the seed in brackets. The lesion summaries do not vary substantially between seeds. Note the x-axis labels denote the center of the age bins, the y-axis units are in 2mm^3^ voxels, and the variance GRF parameter is fixed to 1 for all simulation settings.

**Figure S4:**
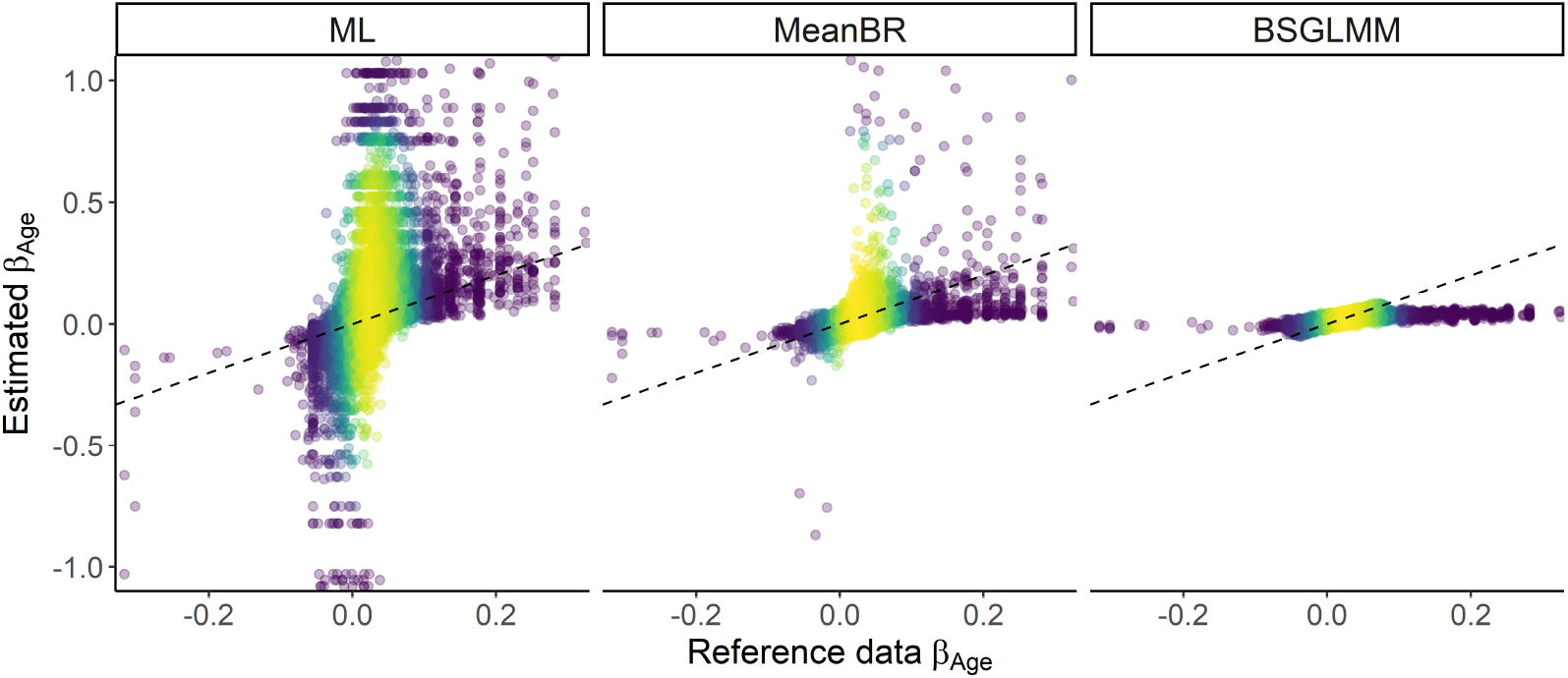
Estimated coefficients 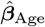 (ML), 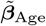 (MeanBR), 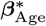 (BSGLMM) vs. ***β***_Age_ (reference). Each point is coloured according to the density of points in an invisible grid overlaid on the plots (the brighter the colour, the higher the density of the points) and the identity superimposed (dashed black line). Bias reduction and the effect of the prior result in shrinkage of the coefficients towards zero with the Bayesian model following the equality line most closely. The ‘horizontal effect’ observed mostly at the BSGLMM plot (826 voxels have reference data coefficients greater than 0.1 in absolute value) occurs when the lesion incidence is very low. One simulated data set of 1,000 subjects used; 40,338 voxels with finite MLEs plotted.

**Figure S5:**
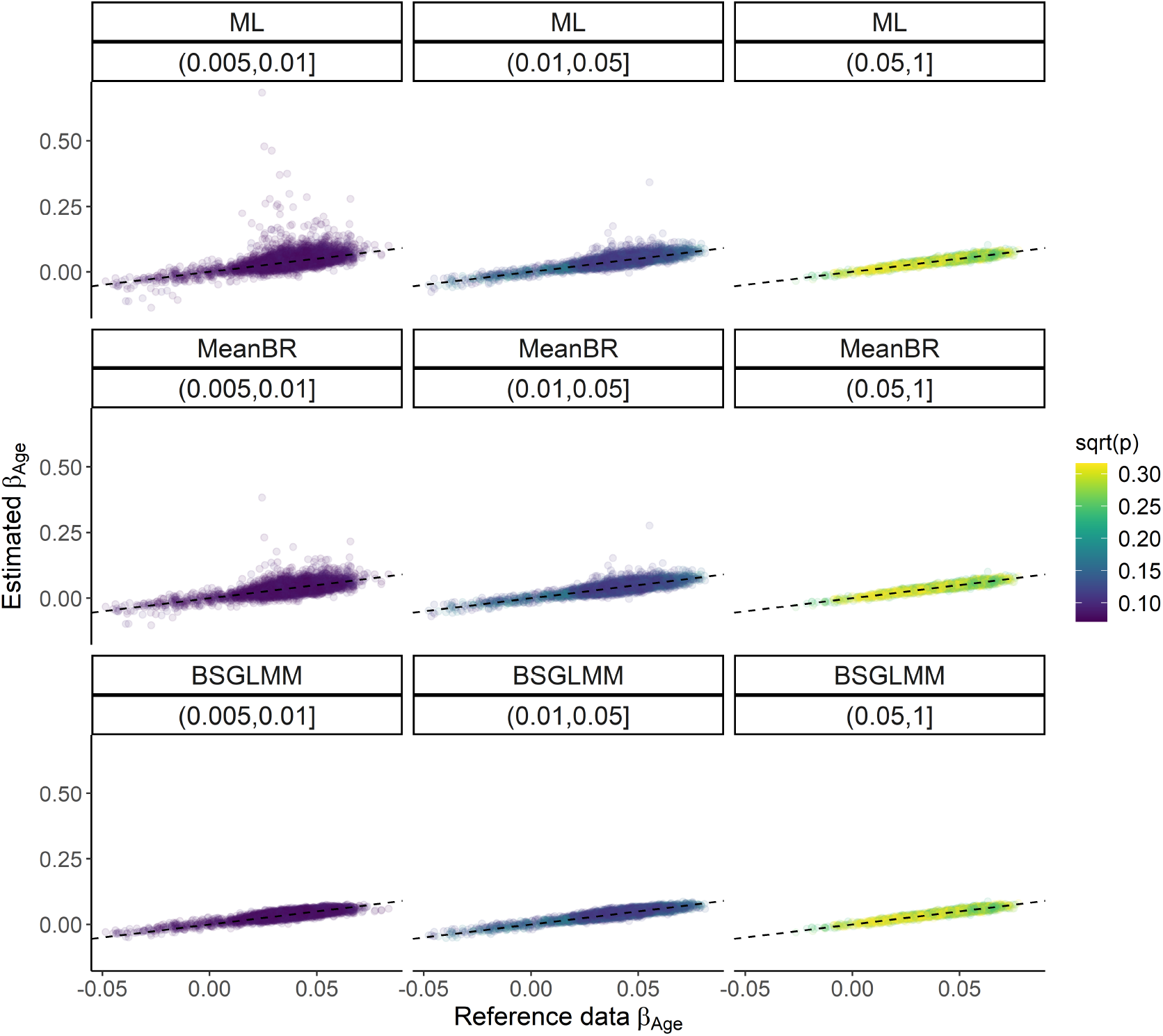
Estimated coefficients 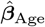 (ML), 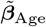 (MeanBR), 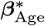 (BSGLMM) vs. ***β***_Age_ (reference) across bins of voxels. Each point is coloured according to the square-root lesion probability 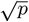 suggesting shrinkage is observed for voxels with low lesion incidence. One simulated data set of 1,000 subjects used; 11,632 voxels with reference data lesion incidence *p* > 0.005 and finite MLEs plotted.

**Figure S6:**
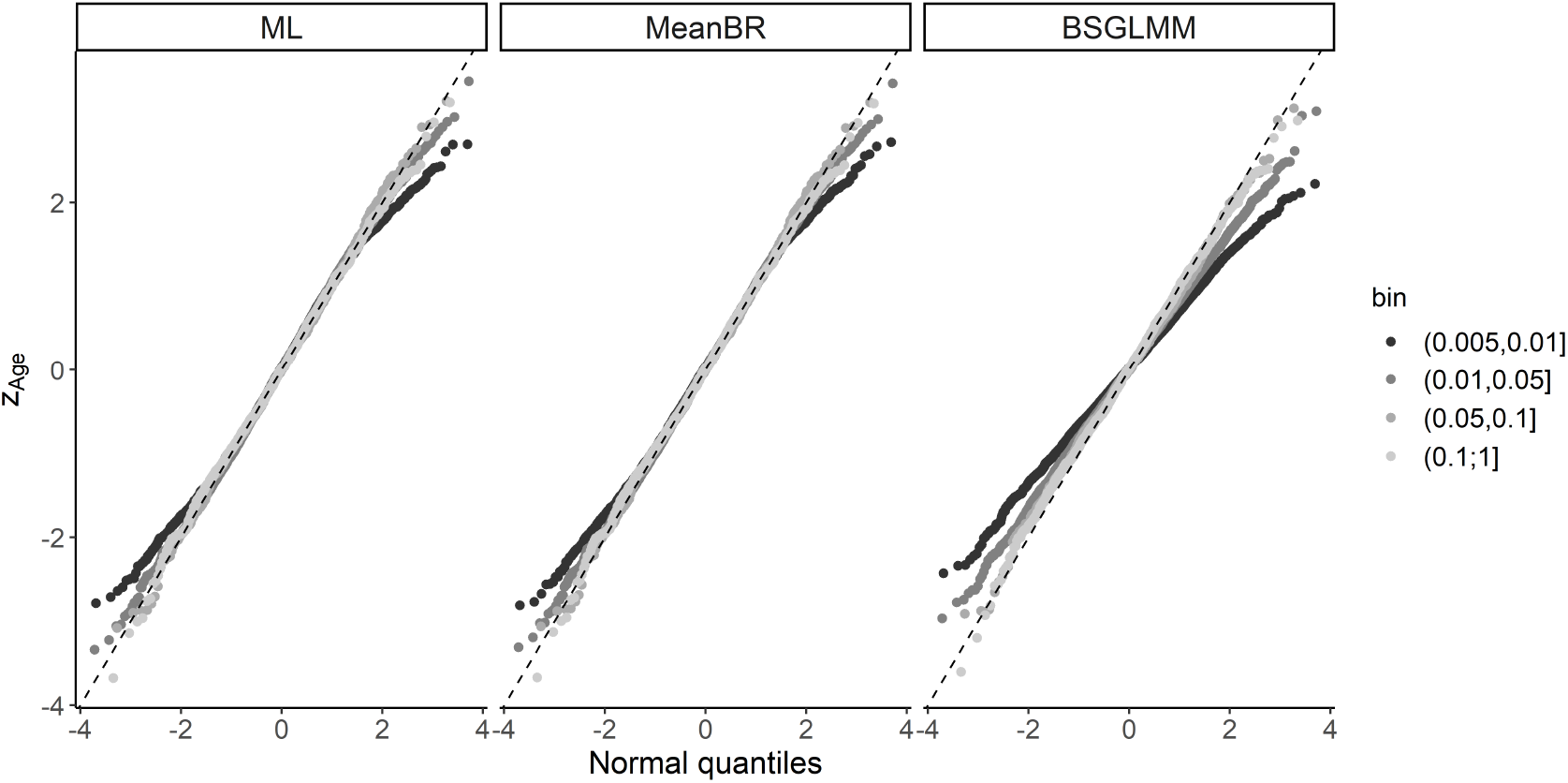
Quantile-quantile (QQ) plots of the quantiles of the simulated data z-scores across bins of voxels versus the theoretical quantiles from a Normal distribution. The lower the lesion incidence (darker colour), the greater the deviations from a linear trend, i.e. the rarer the lesions, the greater the deviations from normality.

**Figure S7:**
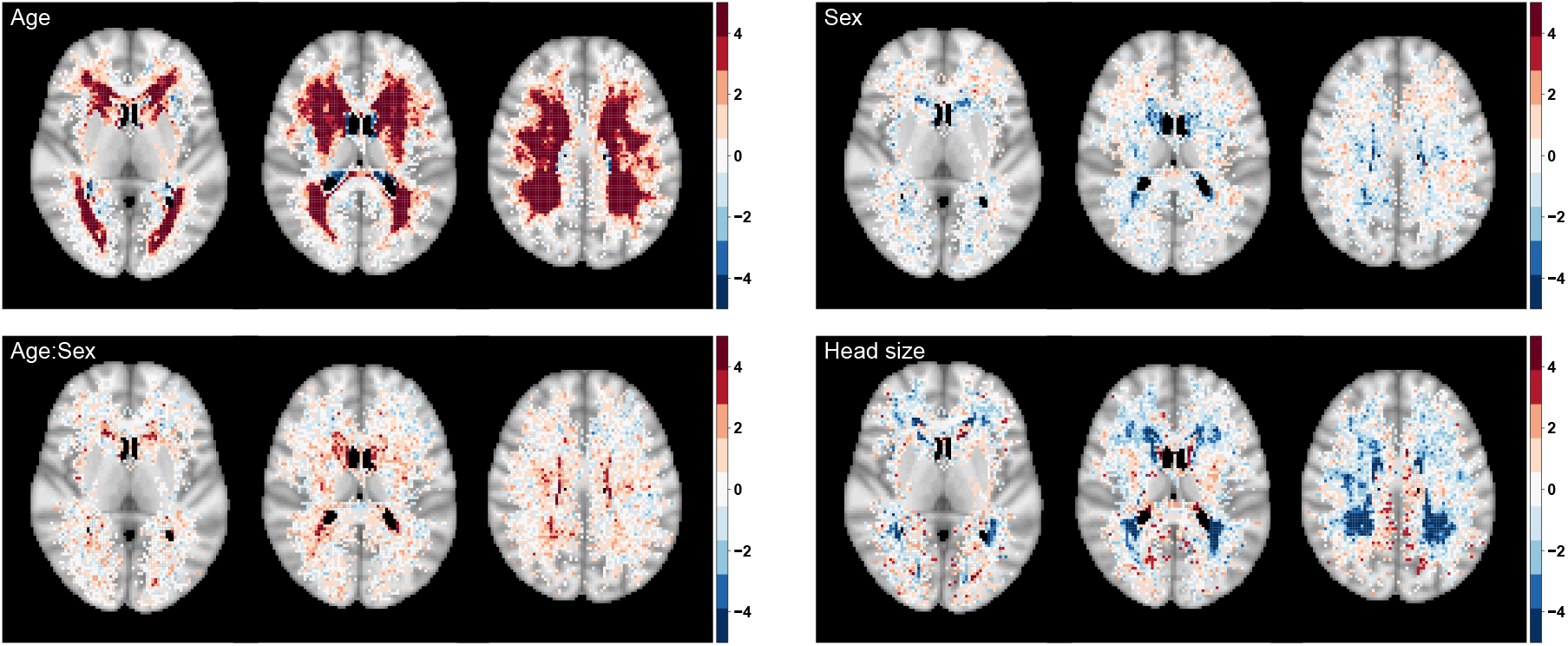
Significance maps (z-scores based on MeanBR estimates) for the effect of age, sex (baseline men), age by sex interaction and head size scaling to complement Figure 5. Data on 13,680 UK Biobank participants used. 72,603 voxels with non-zero lesion probability shown with zero-lesion incidence voxels plotted as transparent to show anatomical MRI for reference; axial slices {40, 45, 50} shown.

## Footnotes

1 Project URL: https://osf.io/h7sxr/

2 Software packages handle separation differently depending on their convergence criterion and the user might not be notified.

3 Code available at https://www.nisox.org/Software/BSGLMM/

4 https://fsl.fmrib.ox.ac.uk/fsl/fslwiki/Cluster

5 Project URL: https://osf.io/h7sxr/

6 Project URL: https://osf.io/h7sxr

